# Diverse Genetic Determinants of Nitrofurantoin Resistance in UK *Escherichia coli*

**DOI:** 10.1101/2021.05.27.446087

**Authors:** Yu Wan, Ewurabena Mills, Rhoda C.Y. Leung, Ana Vieira, Elita Jauneikaite, Xiangyun Zhi, Nicholas J. Croucher, Neil Woodford, Matthew J. Ellington, Shiranee Sriskandan

## Abstract

Antimicrobial resistance in enteric or urinary *Escherichia coli* is a risk factor for invasive *E. coli* infections. Due to widespread trimethoprim resistance amongst urinary *E. coli* and increased bacteraemia incidence, a national recommendation to prescribe nitrofurantoin for uncomplicated urinary tract infection was made in 2018. Nitrofurantoin resistance is reported in <6% urinary *E. coli* isolates in the UK. However, mechanisms underpinning nitrofurantoin resistance in these isolates remain unknown. This study aimed to identify genetic determinants of nitrofurantoin resistance in a local *E. coli* collection and assess their prevalence in a larger dataset of *E. coli* genomes. Deleterious point mutations and gene-inactivating insertion sequences in both chromosomal nitroreductase genes *nfsA* and *nfsB* were identified in genomes of nine nitrofurantoin-resistant urinary *E. coli* isolates collected from north west London. Eight types of genetic alterations were identified when comparing sequences of *nfsA*, *nfsB*, and the associated gene *ribE* in 12,412 *E. coli* genomes collected from across the UK. Evolutionary analysis revealed homoplasic mutations and explained the order of stepwise mutations. An algorithm was developed to predict nitrofurantoin susceptibility and predictions for 20 accessible isolates were experimentally validated. Only one genome carrying *oqxAB*, a mobile gene complex associated with reduced nitrofurantoin susceptibility, was identified. In conclusion, mutations and insertion sequences in *nfsA* and *nfsB* are leading causes of nitrofurantoin resistance in UK *E. coli*. As nitrofurantoin exposure increases in human populations, the prevalence of nitrofurantoin resistance in carriage *E. coli* isolates and those from urinary and bloodstream infections should be monitored.

**Importance:** This study expands knowledge about the genetic basis of nitrofurantoin resistance in *E. coli* isolates using whole-genome sequencing and genomic analysis. We report novel and previously known deleterious mutations of chromosomal genes *nfsA*, *nfsB*, and *ribE* as well as the interruption of *nfsA* and *nfsB* by insertion sequences, recapitulating the roles of oxygen-insensitive nitroreductases in the development of nitrofurantoin resistance in *E. coli*. We revealed and categorised the genotypic diversity in these three genes in a large collection of UK *E. coli* genomes. A scoring algorithm is provided to predict nitrofurantoin susceptibility from genotypes. Our predictions suggest that acquired nitrofurantoin resistance is not of immediate concern in the UK. However, experimental validation of predictions suggested the involvement of mechanisms other than alterations in *nfsA*, *nfsB*, or *ribE* in determining nitrofurantoin susceptibility, emphasising the need for monitoring nitrofurantoin resistance amongst *E. coli*.

## Introduction

Nitrofurantoin is a synthetic nitrofuran compound that has been widely used as a first-line antimicrobial agent for treating urinary tract infection (UTI) since 1953 and is active against a broad range of Gram-negative and Gram-positive bacteria, including *Escherichia coli*, most species of *Staphylococcus* and *Enterococcus* (1, 2). It can reach a urinary concentration of 50–250 mg/L while retaining negligible blood concentration at standard therapeutic dosage, providing an advantage for treating UTI (3, 4). Although the antibacterial activity of nitrofurantoin is not fully understood, studies have revealed that metabolic intermediates from nitroreductase-mediated reduction of nitrofurantoin can damage DNA and RNA, as well as disrupting protein production (5–7).

*E. coli* is the predominant pathogen of uncomplicated UTI, and the increasing prevalence of isolates resistant to antimicrobials such as beta-lactams, ampicillin, and trimethoprim (8, 9) has led the UK National Institute for Health and Care Excellence (NICE, www.nice.org.uk) to recommend prescribing nitrofurantoin for UTI since November 2018 (10). Despite clinical use of nitrofurantoin for nearly 70 years, the prevalence of nitrofurantoin resistance in *E. coli* remains relatively low in Europe. Up to 2016, fewer than 6% of *E. coli* isolates collected from urine specimens in Western, Northern, and Southern European countries were resistant to nitrofurantoin (11–15). Separate UK-based studies showed that nitrofurantoin-resistant *E. coli* accounted for 5% of urinary and bloodstream *E. coli* isolates collected in London, during 2005–2006 and 2011–2015, despite higher prevalence of resistance (>20%) to other commonly prescribed oral antimicrobials in the same isolates (16–18).

Since *E. coli* was not widely exposed to nitrofurantoin in the UK until adoption of the new NICE guideline, it may be too early to detect any trend in the prevalence of nitrofurantoin resistance among UK *E. coli* isolates. The reported low prevalence of nitrofurantoin resistance might be explained by the broad intracellular target range of toxic nitrofurantoin metabolites and a reported fitness cost among nitrofurantoin-resistant mutants (5, 19). Two kinds of genetic determinants of nitrofurantoin resistance have been identified in *Enterobacteriaceae* so far: (a) loss-of-function mutations in chromosomal genes *nfsA*, *nfsB* (encoding oxygen-insensitive nitroreductases NfsA and NfsB, respectively), and *ribE* (encoding 6,7-dimethyl-8-ribityllumazine synthase, involved in the reduction of nitrofurantoin) (6, 7); and (b) acquired gene complex *oqxAB* (encoding a multidrug efflux pump OqxAB) (20). By contrast, bacteria with impaired DNA-repair ability showed increased susceptibility to nitrofurantoin (21, 22).

Understanding the mechanisms underpinning antimicrobial resistance (AMR) is central to predicting the impact of antimicrobial prescribing guidelines. Nevertheless, there is little or no understanding of the basis of nitrofurantoin resistance in UK *E. coli*. This is hindered by the fact that nitrofurantoin susceptibility testing is mostly limited to urinary tract isolates rather than enteric or invasive bloodstream isolates, while a vast array of whole-genome sequencing (WGS) has focussed on bloodstream isolates or isolates with multidrug-resistance phenotypes. We set out to uncover the genetic basis of nitrofurantoin resistance in *E. coli* isolated from UTI patients in north west London. We then screened a large genomic dataset of 12,412 UK *E. coli* isolates for resistance-associated genetic alterations to enhance our understanding of the wider distribution of nitrofurantoin resistance and genetic mechanisms that influence nitrofurantoin’s interaction with *E. coli*.

## Results

### Clinical *E. coli* isolates from north west London

Nitrofurantoin resistance (MIC > 64 mg/L) was confirmed in nine out of 18 *E. coli* UTI isolates that had been reported as nitrofurantoin-resistant by an NHS diagnostic laboratory (supplementary Table S1), following protocols and clinical breakpoints specified by the European Committee on Antimicrobial Susceptibility Testing (EUCAST). Only one isolate was obtained from each patient. Genomic DNA from these nine isolates (IN01–09) was subject to WGS and compared with genomic sequences of *E. coli* with known nitrofurantoin susceptibility.

### Population structure of *E. coli* isolates with known nitrofurantoin susceptibility

To contextualise the isolates IN01–09, genomes of additional 208 *E. coli* isolates with known nitrofurantoin-susceptibility phenotypes were incorporated into analysis (Table S1). Of these, eight isolates were nitrofurantoin-resistant, none of which came from the UK, and 200 isolates were nitrofurantoin-susceptible, 14 of which came from the UK.

Population structure of the 217 isolates (including IN01–09) was interpreted using both multi-locus sequence types (MLSTs) and genomic similarities. In total, 57 *E. coli* sequence types (STs) constituting 19 clonal complexes (CCs, defined by the MLST scheme) were identified (Fig. 1 and Table S1). No novel MLST allele or ST was detected. Isolates IN01–09 were classified into seven STs (Table 1). The 217 *E. coli* isolates were further grouped into 38 genomic clusters using PopPUNK (23), and as expected, these clusters showed a high congruence with CCs (Fig. 1). Notably, isolates IN01 and IN02 only slightly differed in presence-absence of accessory genes by a PopPUNK accessory-genome distance (Jaccard distance) of 2.4 × 10^−4^, although they were collected from different patients with distinct community UTI episodes. Interestingly, while the majority of nitrofurantoin-resistant isolates were sporadically distributed across these CCs (including major clinical lineages such as CC69, CC73, and CC131), clustering of reduced nitrofurantoin susceptibility was seen among USA isolates of CC38 (Fig. 1 and Table S1).

**TABLE 1.**
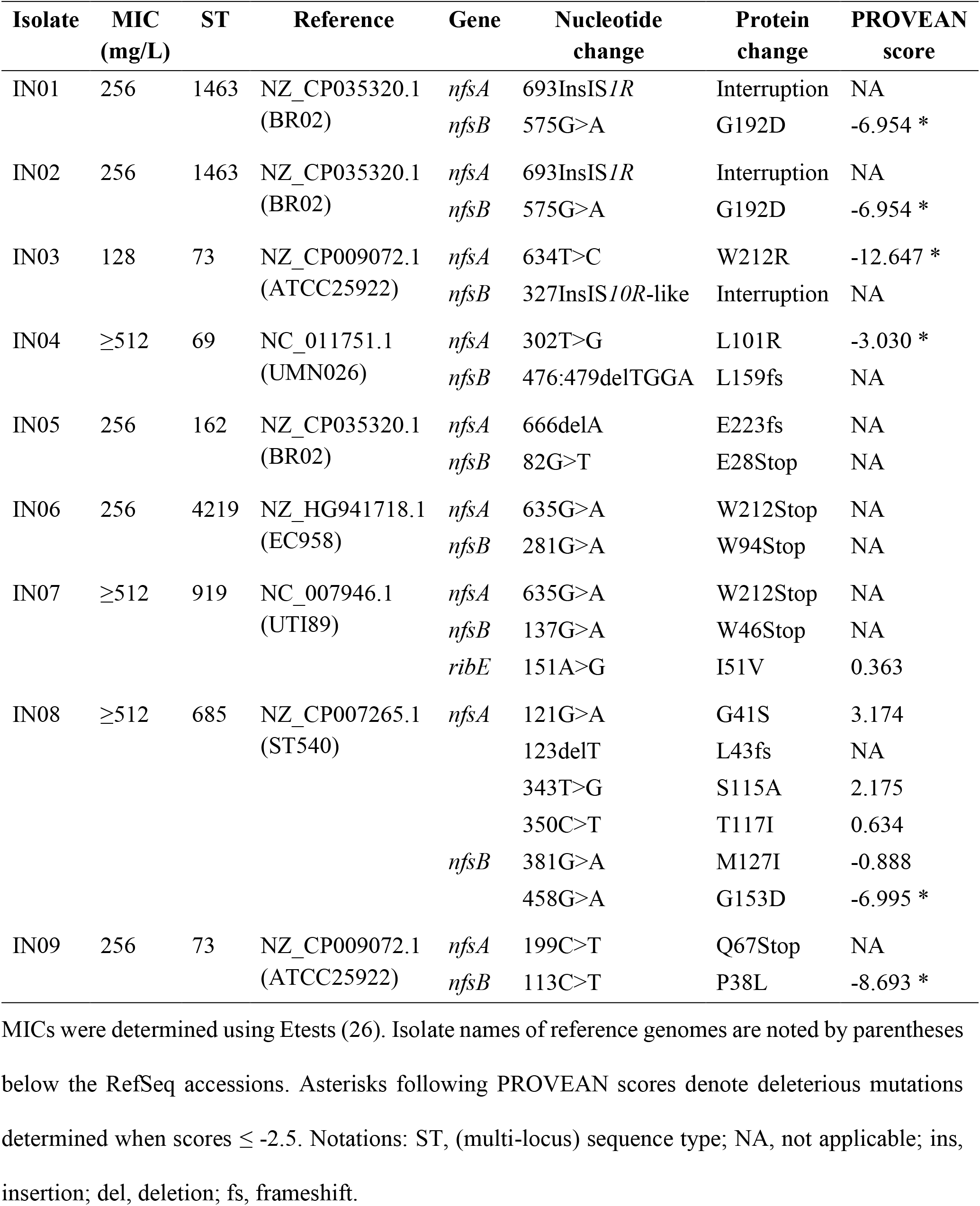
Nonsynonymous mutations and gene interruptions in *nfsA* and *nfsB* of isolates IN01–09.

**FIG 1.**
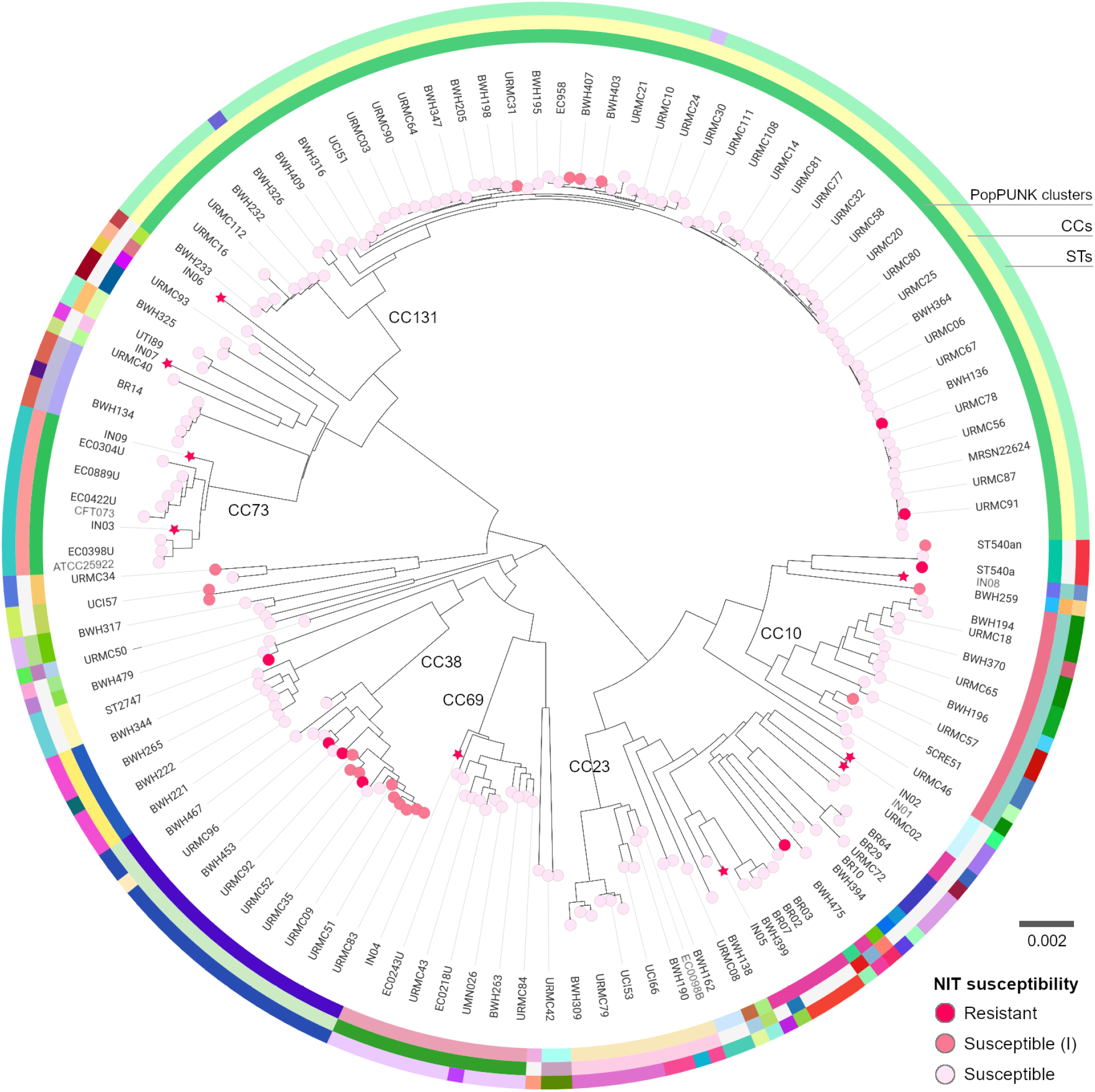
Population structure of 217 *E. coli* isolates with known nitrofurantoin (NIT) susceptibility. Relatedness between these isolates is displayed by a core-genome neighbour-joining tree generated using PopPUNK. The scale bar denotes the density of core-genome single-nucleotide polymorphisms per unit of branch lengths (23). The tree is midpoint-rooted and drawn in a circular layout with tips coloured by nitrofurantoin susceptibility of isolates (Table S1). Stars highlight isolates IN01–09. CCs having ≥10 genomes are labelled over the tree. Following EUCAST guidelines, an isolate was considered susceptible under an increased exposure of nitrofurantoin, denoted as “susceptible (I)”, when 32 mg/L < MIC ≤ 64 mg/L. Full data from this figure can be interactively visualised on Microreact (microreact.org/project/pHwAhGZ55AL7CvXxGnziyU) (24).

### Genetic basis of nitrofurantoin resistance in UK *E. coli* isolates IN01–09

Isolates IN01–09 showed a minimum nitrofurantoin MIC of 128 mg/L, and the MICs of three isolates exceeded 512 mg/L (Table 1). No public WGS data of other nitrofurantoin-resistant UK *E. coli* isolates was found as of May 2020. We focused on alternations in intrinsic genes *nfsA*, *nfsB*, *ribE*, and presence-absence of the acquired gene complex *oqxAB*. For each genome of isolates IN01–09, its most closely related complete genome (based on PopPUNK core-genome distances) of a nitrofurantoin-susceptible *E. coli* isolate was chosen as a reference for the identification of genetic alterations.

### Genetic alterations of nfsA and nfsB

Analysis of *nfsA* and *nfsB* confirmed the presence of a single copy of each gene in each of the nine genomes IN01–09 (Table S2). In each of these genomes, genetic alterations affecting protein sequences were identified in both *nfsA* and *nfsB* when alleles of both genes were compared with those in the reference genome. Altogether, these alterations consisted of missense and nonsense mutations, deletions, and interruptions by insertion sequences (Table 1). Correspondingly, predicted sequences of proteins NfsA and NfsB showed amino acid substitutions or truncations or both. Synonymous mutations in *nfsA* or *nfsB* were also detected in five genomes (Table S2). One deleterious missense mutation was predicted by PROVEAN (25) for each NfsA or NfsB sequence that only carried missense mutations (Table 1). These deleterious missense mutations were not present in any NfsA or NfsB sequence from the other 208 *E. coli* isolates whose nitrofurantoin susceptibility was known. Only two deleterious missense mutations (W212R in NfsA and G192D in NfsB) were previously reported as associated with nitrofurantoin resistance (appendix Tables A1 and A2).

Interruption of *nfsA* or *nfsB* by insertion sequences was identified in three genomes, IN01–03 (Table 1), using SPAdes (27) assembly graphs (Dataset S1) and was then confirmed using Sanger sequencing of PCR products (Fig. 2). Annotations and interrupted genetic regions of these insertion sequences are available at github.com/wanyuac/NITREc/tree/master/Seq/Nucl/IS. Specifically, *nfsA* in both IN01 and IN02 was interrupted by IS*1R* (768 bp, IS*1* family) encoded on the reverse complementary strand. No nucleotide divergence was seen between these two copies of IS*1R* and their reference sequence (GenBank accession: AH003427.2; region: 3,877–4,644). Nine-bp direct repeats (DRs) flanking IS*1R* revealed its insertion site between bases 693 and 694 of *nfsA*. Similarly, *nfsB* in IN03 was interrupted by an IS*10R*-like element (1,329 bp, IS*4*-family) that differed from IS*10R* (GenBank accession: AH003348.2; bases 867–2,195) by 12 nucleotides, consisting of two nucleotide substitutions in the right inverted repeat (IRR) and 10 missense mutations that resulted in four amino acid substitutions in the transposase gene. This element was inserted between bases 327 and 328 of *nfsB*, as indicated by its flanking 9-bp DRs, and in an opposite orientation to *nfsB*. A search of this insertion sequence against GenBank (accessed in August 2020) found two exact matches in *E. coli* genomes of animal and clinical origins, respectively (accessions: CP009578.1 and CU928145.2). A complete Tn*10* variant (GenBank accession: AF162223.1) (28), bounded by an IS*10L*-like element and IS*10R*, was also identified in the assembly graph of IN03. Notably, this IS*10L*-like element only differed from the IS*10R*-like element of IN03 by four nucleotide substitutions in their left inverted repeats (IRLs). Assembly depths and connections of nodes consisting of the IS*10L*-like element, IS*10R*, and Tn*10* variant in the assembly graph of IN03 genome suggested high copy numbers of both insertion sequences and only a single copy of Tn*10*. Taken together, although the transposon did not integrate into *nfsB*, replicative transposition of its IS*10L* or IS*10R* component may cause the interruption of *nfsB*.

**FIG 2.**
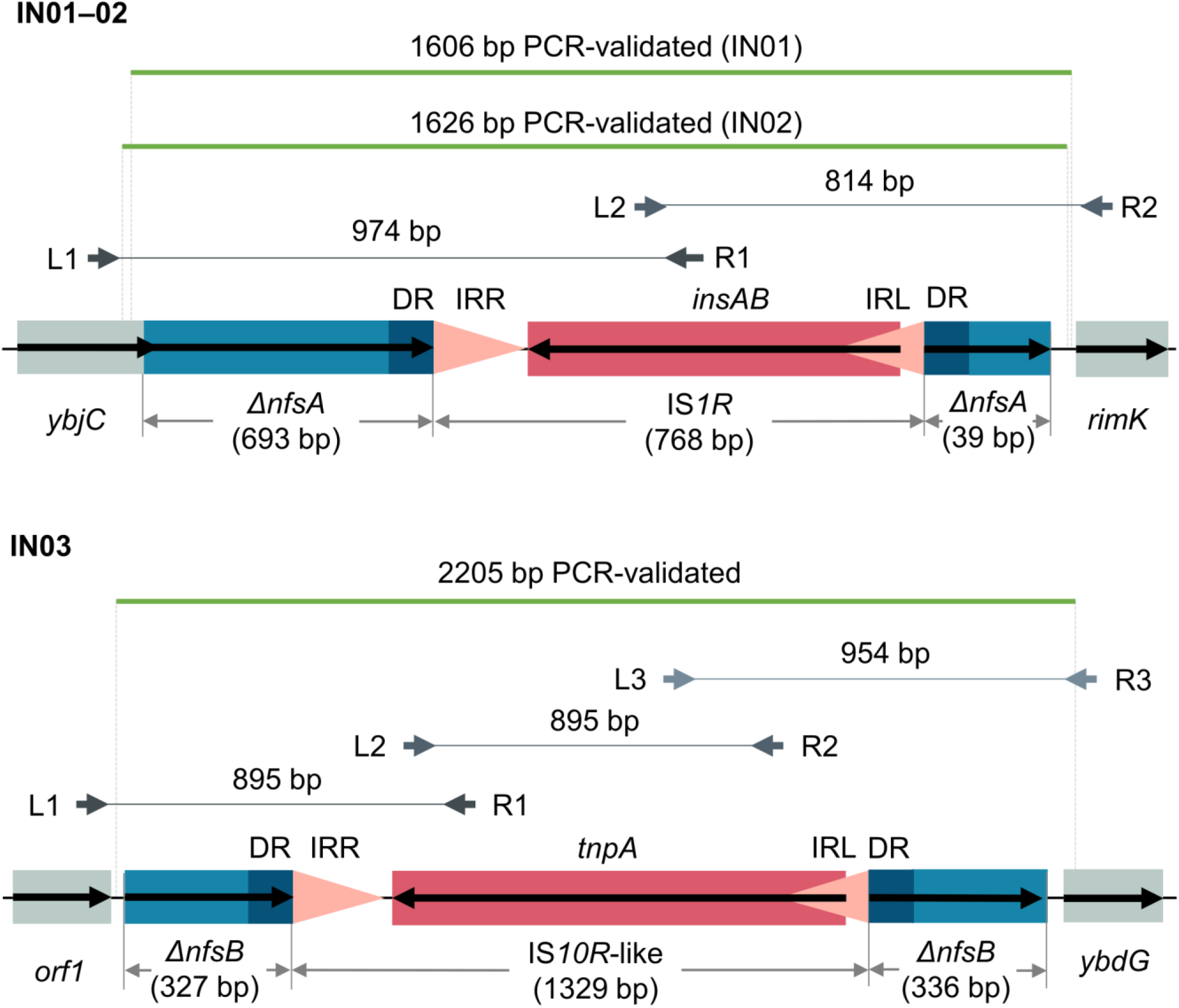
Interruption of *nfsA* and *nfsB* by insertion sequences. Orientation of each coding sequence is indicated by a black bold arrow. IRL and IRR (inverted repeat left and right) of each insertion sequence are represented by pink triangles indicating their orientations. Direct repeats (DRs) flanking each insertion sequence are denoted by dark-blue boxes. These genetic structures were confirmed by Sanger sequencing of PCR products. Paired left and right PCR primers are denoted by grey bold arrows above DNA backbones and are aligned to their source positions. Each pair of primers are connected by a grey solid line to indicate their expected PCR product. Consensus sequences of Sanger reads are denoted by green solid lines. Note that consensus sequences of IN01 and IN02 differed in lengths due to quality trimming of the reads.

### Genetic alterations of ribE

A single copy of *ribE* in each of IN01–09 genomes was predicted from read mapping. Genome IN07 carried the only amino acid substitution identified in RibE (I51V, compared to the reference sequence of genome UTI89), which was attributed to a missense mutation 151A>G (Table S2). Nonetheless, this RibE variant was identical to RibE sequences from another five nitrofurantoin-susceptible reference genomes (ATCC25922, BR02, EC958, ST540, and UMN026). Therefore, the effect of substitution I51V was considered neutral, further supported by PROVEAN prediction (score: 0.363).

### Horizontally acquired nitrofurantoin resistance

No evidence of horizontally acquired nitrofurantoin resistance was identified in isolates IN01–09 as neither the known genetic determinant *oqxA* nor *oqxB* was identified in any of their genomes. Nonetheless, other acquired AMR genes were present in genomes of IN01–05 but not IN06–09 (Tables 2 and S2), conferring resistance to aminoglycosides, beta-lactams, trimethoprim, sulphonamides, and tetracyclines. With one exception (*dfrA1* in IN04), alleles of these genes were identical to the respective reference sequences.

**TABLE 2.**
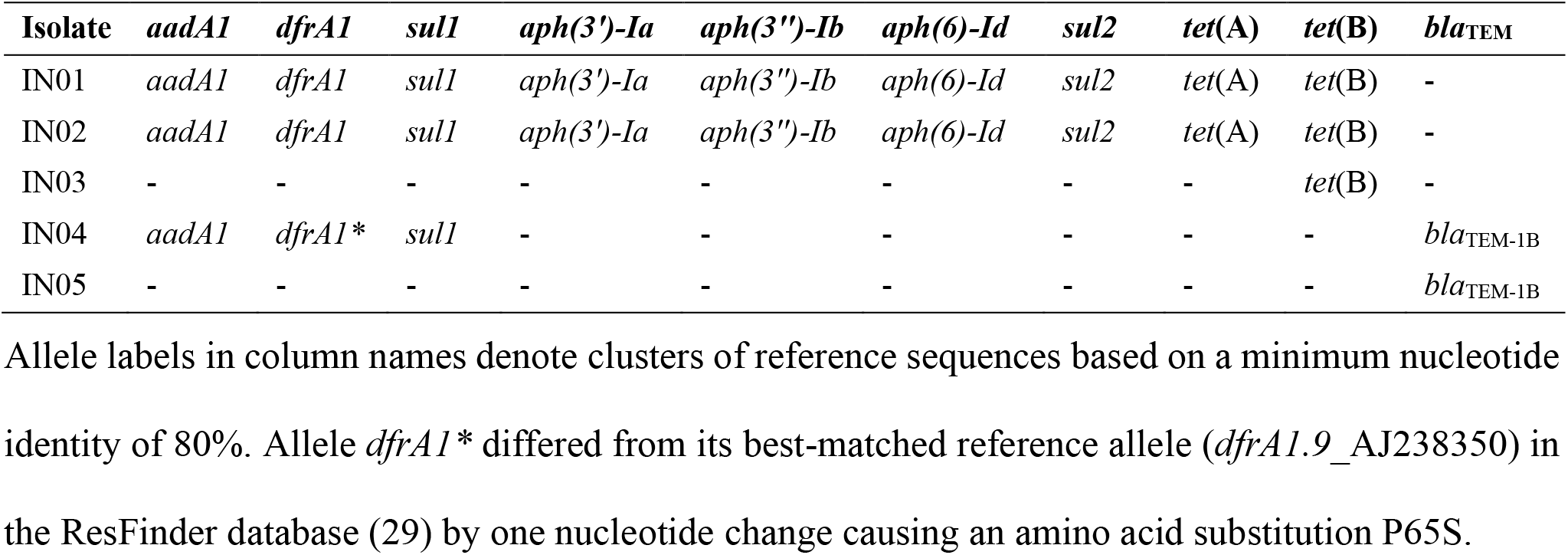
Alleles of acquired AMR genes detected in genomes of isolates IN01–05.

### Diversity of nitrofurantoin-resistance determinants in UK *E. coli*

To contextualise genetic alterations identified in the nine genomes IN01–09, genomes of additional 12,403 UK *E. coli* isolates were retrieved, consisting of 2,009 blood, urine, and gut microbiota isolates from an NIHR Health Protection Research Unit (HPRU) collection; 1,509 blood isolates collected by the British Society for Antimicrobial Chemotherapy (BSAC) or the Cambridge University Hospitals NHS Foundation Trust (hereafter, BSAC or CUH collection) (30); 297 isolates from East England (31); 162 isolates from Scotland (hereafter, SCOT collection) (32); 85 isolates from the NCTC 3000 project (33), including historic isolates; and 8,341 isolates recorded in EnteroBase (34) (Table S3).

Of the 12,412 *E. coli* genomes analysed, nucleotide BLAST identified 12,488 hits for *nfsA* in 12,410 genomes; 12,393 hits for *nfsB* in 12,388 genomes; and 12,417 hits for *ribE* in all 12,412 genomes. Notably, *nfsA* was not detected in two genomes, and *nfsB* was not detected in another 24 genomes. By contrast, 75 genomes (including IN01–02) had 2–4 hits for *nfsA* (complete or partial coding sequence); five genomes (including IN03) had two hits for *nfsB* (partial); and five genomes had two hits for *ribE* (complete). However, none of the 12,412 genomes had more than one hit for any two or all of the genes *nfsA*, *nfsB*, and *ribE*.

A decision tree was developed to categorise hits for each gene based on predicted protein sequences (Fig. 3). Specifically, 164 partial sequences (39–714 bp) of *nfsA* (wildtype length: 723 bp) were identified, which failed to encode complete 240-aa (amino acid) NfsA proteins. Frameshift mutations were present in all 107 alleles with indels, resulting in predicted proteins of 18–247 aa. Nonsense mutations identified in 117 alleles encoded partial NfsA proteins of 23-234 aa. Interestingly, amongst missense mutations, the start codon ATG of *nfsA* was altered in 14 genomes: mutation M1T (2T>C) was found in an allele shared by two BSAC isolates eo393 and eo2899 (collected in 2002 and 2010, respectively), causing a loss of start codon; start-codon substitution M1I (3G>A) was found in five *nfsA* alleles from 12 genomes (Table S4) and was considered deleterious to protein structure according to the PROVEAN prediction (uniform score: −3.149).

**FIG 3.**
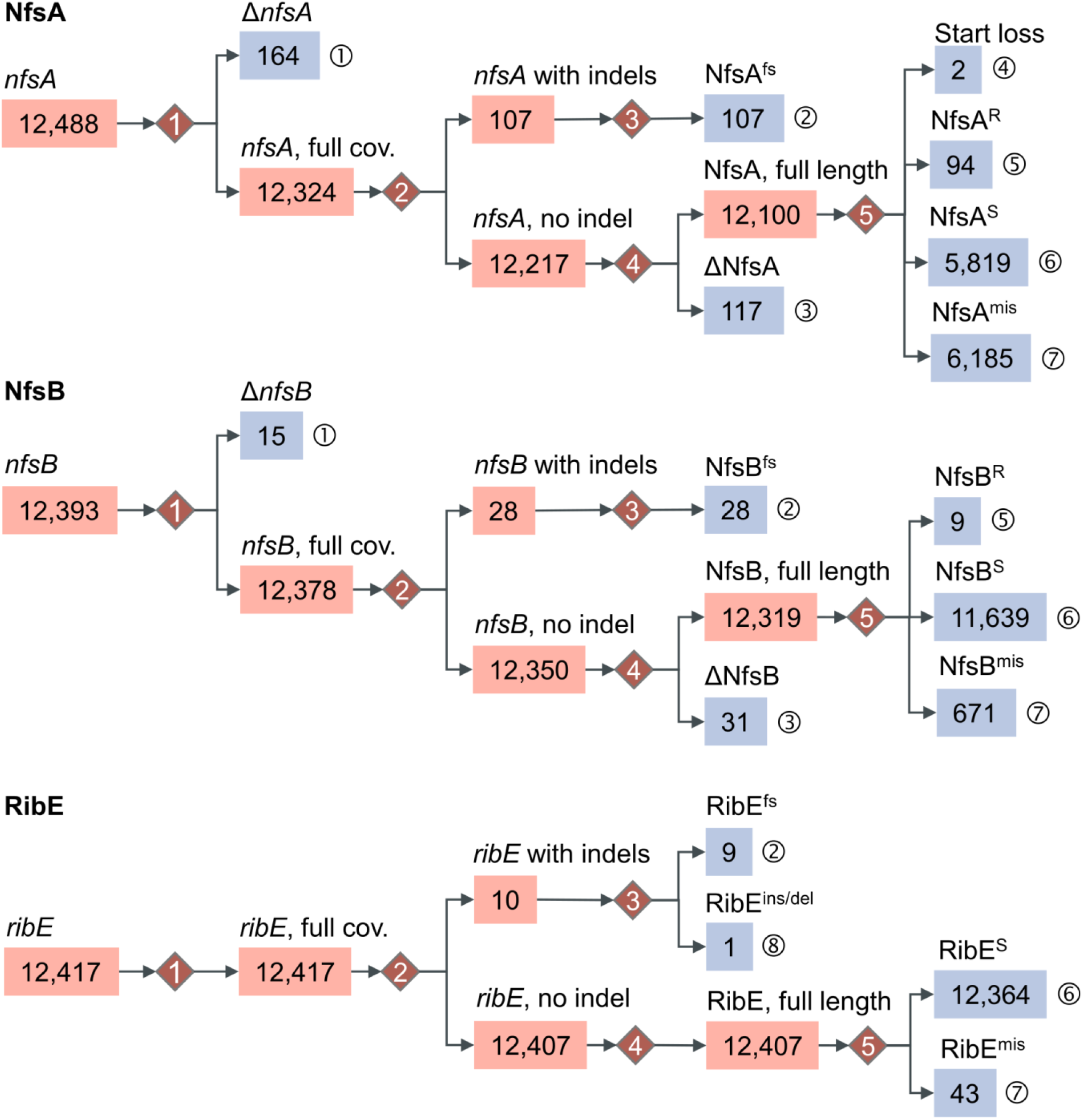
Classification of *nfsA*, *nfsB*, and *ribE* alleles from 12,412 *E. coli* genomes, based on nucleotide- and protein-level comparisons. Final allele categories are labelled with ① – ⑧ and shaded in blue. These categories are determined at five decision points indicated by numbered red diamonds. Numbers of alleles in each category are displayed in boxes. Categories with zero counts are omitted for convenience. Assessment at each decision point: (1) Does the allele sequence cover 100% of its reference sequence? (2) Is there any gap opened in the sequence alignment? (3) Is there a frameshift mutation or insertion/deletion of any codon? (4) Is there any nonsense mutation? (5) Does the predicted protein sequence carry no mutation, or any known resistance-associated amino acid substitution, or any other amino acid substitution? Notations: Δ, partial sequence compared to its reference; cov., query coverage; fs, frameshift mutation; ins/del, insertion or deletion of amino acid residues; R, missense mutations known to be associated with nitrofurantoin resistance; S, proteins identical to those in nitrofurantoin susceptible isolates; mis, missense mutations of unknown phenotypical impacts.

Genetic alterations of *nfsB* (654 bp) displayed a similar pattern to *nfsA*. Particularly, 15 partial sequences (35–647 bp) were identified and no start-codon loss was seen amongst missense mutations (Table S4). By contrast, *ribE* (471 bp) displayed lower diversity than the other two genes, with no partial allele sequence or nonsense mutation identified. At least one copy of *ribE* was detected in each genome.

Among *ribE* alleles identified in the 12,412 genomes, a stop-codon substitution (TGA>TAA) owing to missense mutation 470G>A was found in two *ribE* alleles that were present in 31 genomes. Both alleles encoded the same RibE protein as the nitrofurantoin-susceptible strain ATCC25922 (NCBI protein accession: WP_001021161.1). A novel deletion of four amino acid residues (KAGN, from position 132 to 135 in the reference RibE sequence from the ATCC25922 genome) was predicted from one (carried by isolate EC0430U) of 10 *ribE* alleles that possessed indels. Notably, this deletion overlapped the deletion of amino acid residues 131–134 (TKAG, from the same reference RibE sequence) that is known to reduce nitrofurantoin susceptibility (7). Putatively deleterious missense mutations of RibE were identified in three genomes of further HPRU isolates: EC0340B (A16V), EC0444B (A34T), and EC1165B (T131S), with PROVEAN scores of −3.406, −3.893, and −2.742, respectively (Table S4).

Acquired resistance genes *oqxA* (1,176 bp) and *oqxB* (3,153 bp) were rare among the 12,412 genomes: both genes were detected in only one genome (eo1692, from the BSAC collection), with exact matches to their reference sequences in the ResFinder database (29). No other hit for either gene was obtained given minimum nucleotide coverage and query coverage of 70% and 80%, respectively.

### Evolution of *nfsA*, *nfsB*, and *ribE*

Based on the observed diversity of these three genes in the 12,412 *E. coli* genomes, we hypothesised that *ribE* was under stronger negative or purifying selection than *nfsA* and *nfsB*. The estimated ratio for the nucleotide nonsynonymous-substitution rate (dN) over synonymous-substitution rate (dS), denoted as *ω* = *dN/dS*, was 0.607812, 0.40739, and 0.179068 for *nfsA*, *nfsB*, and *ribE*, respectively, indicating that all three genes were under purifying selection with ascending strength in the order *nfsA* < *nfsB* < *ribE*.

After deduplicating identical alleles from the 12,412 genomes, 231 (32% of 723 bp), 184 (28% of 654 bp), and 86 (18% of 471 bp) SNP sites were found in 328 *nfsA* alleles, 231 *nfsB* alleles, and 88 *ribE* alleles, respectively (Figures S1–3). Despite this variation, phylogenetic trees of these three genes generally showed extremely low bootstrap values (for instance, 0–2) in the majority of ancestral branches (Dataset S1), suggesting a lack of consensus phylogenetic information in these alleles for reliably reconstructing the evolutionary history of each gene.

Pairwise homoplasy indexes (PHIs) calculated from the allele alignment of each gene revealed significant homoplasy in *nfsA* (p-value: 0.0434) and *nfsB* (p-value: 0.0363) but not in *ribE* (p-value: 0.736). Comparisons of each allele alignment to its corresponding gene tree identified 82, 50, and 12 homoplasic SNP sites in *nfsA*, *nfsB*, and *ribE*, respectively (Figures S1–3).

### Prediction of nitrofurantoin susceptibility and experimental validation

For each of the 2,009 HPRU isolates, which we had access to, nitrofurantoin susceptibility was predicted from sequence categories (Fig. 3) of intrinsic genes *nfsA*, *nfsB*, *ribE* and presence-absence of both acquired genes *oqxA* and *oqxB* using a scoring algorithm (Section “Materials and Methods”). Specifically, the algorithm gave a risk score *r* (*0* ≤ *r* ≤ *4*) to each isolate as a summary of its five-gene (*nfsA*, *nfsB*, *ribE*, *oqxA*, and *oqxB*) sequence categories or genotypes. We hypothesised that a higher score would predict a greater reduction in the nitrofurantoin susceptibility of an isolate, and vice versa. Of note, neither *oqxA* nor functional *oqxB* was detected in any of these isolates.

Using *r* ≥ *1* as a criterion, a non-redundant set of 62 HPRU *E. coli* isolates (one isolate per patient) were predicted to have reduced nitrofurantoin susceptibility (Table S5). Since RibE sequences of these isolates were identical to those of wildtype RibE proteins in nitrofurantoin-susceptible isolates, 16 genotypes associated with reduced nitrofurantoin susceptibility were identified in these 62 isolates based on *nfsA* and *nfsB* sequences only (Table 3).

**TABLE 3.** *nfsA-nfsB* genotypes of *E. coli* isolates with nitrofurantoin-resistance risk scores ≥ 1.

|  |  | Isolate number of each genotype (number selected for experimental validation) |  |  |  |  |  |
| --- | --- | --- | --- | --- | --- | --- | --- |
| *nfsA* \ *nfsB* | Wildtype | Missense, known | Missense, unknown | Nonsense | Frameshift | Fragmented | Absent |
| Wildtype | 0 | 1 (1) | 0 | 0 | 1 (1) | 0 | 0 |
| Missense, known | 9 (3) | 0 | 3 (2) | 1 (1) | 0 | 0 | 1 (1) |
| Missense, unknown | 0 | 0 | 0 | 0 | 2 (1) | 0 | 1 (1) |
| Nonsense | 13 (1) | 1 (1) | 1 (1) | 0 | 1 (1) | 0 | 0 |
| Frameshift | 13 (1) | 0 | 0 | 0 | 0 | 0 | 0 |
| Fragmented | 11 (1) | 1 (1) | 2 (2) | 0 | 0 | 0 | 0 |

Genotypes of 62 non-redundant HPRU *E. coli* isolates are shown. A gene was considered fragmented when it was interrupted or truncated. In total, 16 genotypes had at least one isolate each, and 20 isolates were selected for experimental validation.

Assuming nonsense or frameshift mutations observed in this study always cause a loss of function, twenty representative isolates of the 16 genotypes were selected from the 62 isolates as a “prediction group” for quantitative antimicrobial susceptibility testing (Table 4). Specifically, one representative was chosen for each genotype that did not involve any missense mutation. Otherwise, one representative was chosen for each missense mutation. In this group, all 12 double mutants carried at least one resistance-associated genetic alteration (*1* < *r* ≤ *2*). Nitrofurantoin susceptibility was correctly predicted for all these isolates. Notably, the double mutants were highly resistant to nitrofurantoin (MIC ≥ 256 mg/L) when both genetic alterations were known to be associated with nitrofurantoin resistance. Predictions for the eight single mutants of resistance-associated genetic alterations (*r* = *1*) were less reliable than those for double mutants, with five single mutants being more susceptible to nitrofurantoin than predicted. Isolates EC394_9, EC0880B, and EC0026B showed heterogenous responses to nitrofurantoin as a few colonies of resistant *E. coli* appeared within inhibition zones.

**TABLE 4.** Genotypes and confirmed nitrofurantoin susceptibility of *E. coli* isolates in test groups.

| Group | Isolate | NfsA<br>(240 aa) | NfsB<br>(217 aa) | RibE<br>(156 aa) | *r* | Susceptibility |  | MIC<br>(mg/L) |
| --- | --- | --- | --- | --- | --- | --- | --- | --- |
|  |  |  |  |  |  | Predicted | Observed |  |
| Prediction<br>(20 isolates) | EC0064B | G154E | Wildtype | Wildtype | 1.0 | S(I) | S(I) | 64 |
|  | EC0890B | H11Y | Wildtype | Wildtype | 1.0 | S(I) | S | 16 |
|  | EC1069B_1 | G126R | Wildtype | Wildtype | 1.0 | S(I) | S | 32 |
|  | EC394_9 | Q67Stop | Wildtype | Wildtype | 1.0 | S(I) | S(I) | 64 * |
|  | EC0880B | 1:421int413:723 | Wildtype | Wildtype | 1.0 | S(I) | R | 256 * |
|  | EC1161B | I228fs | Wildtype | Wildtype | 1.0 | S(I) | S | 32 |
|  | EC0932B | Wildtype | F84S | Wildtype | 1.0 | S(I) | S | 4 |
|  | EC1187B | Wildtype | Y183fs | Wildtype | 1.0 | S(I) | S | 32 |
|  | EC0026B | Q113Stop | H80Y | Wildtype | 1.1 | S(I) or R | S(I) | 64 * |
|  | EC0067B | 1:308int434:723 | W46R | Wildtype | 1.1 | S(I) or R | R | >512 |
|  | EC0179B | G126R | W94Stop | Wildtype | 2.0 | R | R | >512 |
|  | EC0328B | G126R | Absent | Wildtype | 2.0 | R | R | 256 |
|  | EC0363B | R133C | T88fs | Wildtype | 1.1 | S(I) or R | R | 128 |
|  | EC0439B | H11Y | W46R | Wildtype | 1.1 | S(I) or R | R | >512 |
|  | EC0553Bo | S38F | Absent | Wildtype | 1.1 | S(I) or R | R | >512 |
|  | EC0629B | 1:52del | G192S | Wildtype | 2.0 | R | R | 512 |
|  | EC0644B | 1:69del | P209L | Wildtype | 1.1 | S(I) or R | S(I) | 64 |
|  | EC0812U | W159Stop | L22fs | Wildtype | 2.0 | R | R | >512 |
|  | EC0856B | G131D | K205E | Wildtype | 1.1 | S(I) or R | R | >512 |
|  | EC1131B | E75Stop | G192D | Wildtype | 2.0 | R | R | >512 |
| Exploration<br>(7 isolates) | EC6002_8 | G41S | Wildtype | Wildtype | 0.1 | S or S(I) | S | 8 |
|  | EC1146B | G66R | Wildtype | Alt. stop | 0.1 | S or S(I) | S | 32 |
|  | EC6125_8 | Wildtype | M127I | Wildtype | 0.1 | S or S(I) | S | 16 |
|  | EC0340B | Wildtype | Wildtype | A16V | 0.1 | S or S(I) | S | 16 |
|  | EC0430U | Wildtype | Wildtype | 132:135del | 0.1 | S or S(I) | S(I) | 64 |
|  | EC0444B | Wildtype | Wildtype | A34T | 0.1 | S or S(I) | S | 32 |
|  | EC1165B | Wildtype | Wildtype | T131S | 0.1 | S or S(I) | S | 8 |
| Control (-) | ATCC25922 | Wildtype | Wildtype | Wildtype | 0.0 | S | S | 16 |
| Control (+) | *IN09* | **Q67Stop** | P38L | Wildtype | 1.1 | S(I) or R | R | 256 |

Lengths of wildtype NfsA, NfsB, and RibE are noted beneath protein names. Names of isolates carrying two mutations (namely, double mutants) are italicised. Known genetic alterations associated with nitrofurantoin resistance (Tables 1, A1, and A2) are shaded in red. Genetic alterations identified in IN01–09 genomes are boldfaced. Asterisks indicate isolates of which colonies within inhibition zones were seen and considered when determining the MICs. Interpretation of MICs: S, susceptible ( $\text{MIC} \leq 32 \text{ mg/L}$ ); S(I), susceptible, under increased exposure to nitrofurantoin ( $32 \text{ mg/L} < \text{MIC} \leq 64 \text{ mg/L}$ ); R, resistant ( $\text{MIC} > 64 \text{ mg/L}$ ). Nitrofurantoin susceptibility of the negative (-) and positive (+) controls must not be used as support for validation because their *nfsA*, *nfsB*, and *ribE* alleles were used as references by the prediction algorithm. Abbreviations: int, gene interruption, with nucleotide positions (start : end) of remnants related to their wildtype references noted aside; del, deletion or truncation, with its start and end nucleotide positions noted aside; alt. stop, alternative stop codon.

Seven additional isolates were included as an exploratory group to examine potential effects of novel mutations on nitrofurantoin susceptibility (Table 4): NfsA or NfsB of two isolates had a missense mutation that had also been identified in isolate IN08; *ribE* of one isolate was terminated by an alternative stop codon TAA; and another four isolates had mutations in RibE, while genomes of all the four isolates encoded wildtype NfsA and NfsB. A nitrofurantoin-resistance risk score of 0.1 was assigned to each isolate by the prediction algorithm to denote presence of a novel mutation. All seven isolates were susceptible to nitrofurantoin (MICs ≤ 64 mg/L), matching our predictions. None of the six missense mutations resulted in nitrofurantoin resistance. By contrast, isolate EC0430U, which carried a deletion of KAGN^132–135^ in RibE, showed an MIC of 64 mg/L, exceeding that of a previously reported isolate having an overlapping 4-aa deletion in RibE (7).

### Predicted occurrences of reduced nitrofurantoin susceptibility in UK E. coli blood isolates

As data relating to nitrofurantoin susceptibility among bacteriaemia *E. coli* isolates is seldom available, occurrences of reduced nitrofurantoin susceptibility were calculated for a non-redundant collection of 2,253 bacteriaemia *E. coli* isolates that were collected in 2001–2016 from across the UK, predating the change in national guidelines. This collection comprised 582 HPRU isolates (one isolate was selected per ST per patient), 1,509 BSAC or CUH isolates, and 162 SCOT isolates. Overall, 142 (6.3%) isolates were predicted to show reduced nitrofurantoin susceptibility. Specifically, 123 (5.5%) single-mutation isolates had a resistance-associated alteration in either *nfsA* (117 isolates, 5.2%) or *nfsB* (6 isolates, 0.3%) and 19 (0.8%) double-mutation isolates had resistance-associated alterations in both *nfsA* and *nfsB*.

## Discussion

In this study, we have explored genetic determinants of nitrofurantoin resistance in UK *E. coli* starting with nine sequenced nitrofurantoin-resistant clinical isolates and a comparative genomics approach. We found four types of alterations in two intrinsic, oxygen-insensitive nitroreductase genes (*nfsA* and *nfsB*) that are known to be associated with nitrofurantoin resistance: gene interruptions by insertion sequences, frameshift mutations caused by indels, nonsense mutations and missense mutations both caused by single-nucleotide substitutions. Notably, each of these nine isolates had alterations in both *nfsA* and *nfsB*, equivalent to double-step or multi-step mutants of both genes, and hence explaining high nitrofurantoin MICs (≥ 128 mg/L) of these isolates (35).

Both *nfsA* and *nfsB* had at most one missense mutation identified in genomes of isolates IN01–07 and IN09, and these mutations were predicted to be deleterious to protein functions by PROVEAN (Table 1). Mutation W212R of NfsA in isolate IN03 was previously identified in an Iranian nitrofurantoin-resistant *E. coli* isolate EC168 (MIC ≥ 512 mg/L), in which no functional *nfsB* gene was detected (36). Since nucleotide substitutions differ between IN03 (634T>C) and EC168 (634T>A), this mutation may have arisen independently in each isolate. W^212^, located in α-helix α10, forms the active site of NfsA (Protein Data Bank or PDB accession for protein structure: 1F5V) with the hydrophobic side chain of tryptophan and is conserved when compared with the NfsA counterparts in *Vibrio harveyi* and *Bacillus subtilus* (37, 38). A functional disruption can hence be anticipated when substituting this side chain with the positively charged side chain of arginine. The other missense mutation of NfsA identified in our study, L101R in isolate IN04, is novel and, as in the case of EC168, this mutation possibly disrupts the function of NfsA because IN04 was highly resistant to nitrofurantoin (MIC ≥ 512 mg/L) and did not possess a functional *nfsB* gene. Since L^101^ is part of α-helix α6 in the central domain of NfsA (37, 38), substitution of this hydrophobic, non-polar residue with arginine may have a disruptive impact on the protein structure and function.

Mutation G192D in NfsB from isolates IN01 and IN02 was previously identified in a nitrofurantoin-resistant *E. coli* isolate (collected in 1999–2000) having a deletion in *nfsA* and showing a nitrofurantoin MIC of 128 mg/L (35, 39); substitution of G^192^ with alanine has also been associated with nitrofurantoin resistance (35). Located at the end of the fourth β-sheet of NfsB (PDB accession: 1DS7), G^192^ constitutes the hydrophobic core that accommodates co-factor flavin mononucleotide (FMN) (40) and is in close spatial proximity of negatively charged residue D^160^. Therefore, substitution of this neutral, hydrophobic residue with another aspartic acid residue possibly affects the protein formation as well as its function. A similar impact of the NfsB mutation G153D in isolate IN08 on the protein function is anticipated based on the same difference in hydrophobicity of amino acid residues.

Nonsense mutations, frameshift mutations, and insertion sequence mediated interruptions of *nfsA* or *nfsB* are also common loss-of-function genetic alterations causing nitrofurantoin resistance (35, 36, 41) and we found examples of all these alterations in isolates IN01–09 (Table 1). The interruption of *nfsA* by an IS*1*-family insertion sequence has been reported by three other studies, two of which also identified insertion of IS*1* in *nfsB*, while one identified integration of the composite transposon Tn*10* into *nfsA* (6, 7, 35). Interestingly, the position and flanking nucleotides of the IS*1* insertion site differ among all three studies, indicating variability of IS*1* in gene inactivation and being consistent with the known AT-rich specificity of IS*1* for target sites (42). In comparison, the interruption of *nfsB* by a novel IS*10R*-like insertion sequence had not been reported before, although another IS*4*-family insertion sequence (IS*186*) was found in disrupted *nfsA* (6). Since IS*10*-group elements comprise both ends of Tn*10*, the presence of Tn*10* in isolate IN03 implies transposition of IS*10R* or its IS*10L*-like element from Tn*10* to *nfsA* or *nfsB*, conferring reduced nitrofurantoin susceptibility to host bacteria. Nevertheless, we anticipate that it is less likely for IS*10L* (left end of Tn*10*) to show the same behaviour as IS*10R* because its transposition function is much weaker than the latter (43). We might see a growing frequency of insertion sequence mediated nitrofurantoin resistance in the future, if copy numbers of IS*1*- and *IS4*-family elements in *E. coli* genomes are undergoing similar accumulation trajectories as those in *Shigella* spp (44).

This study has revealed variation of genes *nfsA*, *nfsB*, and *ribE* among UK *E. coli* isolates. Identified alleles of these genes (Fig. 3) can be summarised into two primary types: functional alleles and pseudogenes. The latter can result from gene truncation, insertion sequence mediated gene interruption, frameshift mutations, nonsense mutations, as well as deleterious missense mutations, and confers reduced nitrofurantoin susceptibility to *E. coli*. Comparisons between dN/dS ratios of *nfsA*, *nfsB*, and *ribE* support our hypothesis that *ribE* is subjected to the strongest pressure of negative selection whereas *nfsA* is subjected to the weakest. Not only does this variation in selective pressures explain the observed diversity difference between these three genes in both this study and literature (Tables A1–3), but also explains why *nfsA* usually yields first-step mutants in the scenario of stepwise increments of nitrofurantoin-resistance levels, which has been reported by several studies (6, 7, 35). Notably, *nfsA* and *nfsB* are classified as non-essential genes of *E. coli*, whereas *ribE* is considered as an essential gene by the PEC database (shigen.nig.ac.jp/ecoli/pec) (45). The higher level of evolutionary conservation of *ribE* compared with *nfsA* and *nfsB* appears to represent a greater degree of functional constraint on its sequence (46). Nonetheless, expression levels of these three genes might also be important to consider (47).

Homoplasic SNPs were identified in *nfsA*, *nfsB*, and *ribE* of *E. coli* isolates in our collection (Figures S1–3), and recombination within *nfsA* and *nfsB* was also inferred by the pairwise homoplasy indexes. Both factors may contribute to the lack of congruent phylogenetic signals in alleles of each gene (represented as low bootstrap values of branches in each gene tree), indicating that, in addition to recombination, resistance mutations can arise independently in each gene across the *E. coli* population. Given the observed extremely low occurrence of acquired resistance gene complex *oqxAB* in our *E. coli* collection, *de novo* mutations of chromosomal genes *nfsA*, *nfsB*, and *ribE*; and insertion sequence mediated interruptions of *nfsA* and *nfsB* may constitute the main source of reduced nitrofurantoin susceptibility in UK *E. coli* isolates.

We developed a decision-tree based algorithm for predicting nitrofurantoin susceptibility from five loci (*nfsA*, *nfsB*, *ribE*, *oxqA*, and *oqxB*) that are known to be involved in nitrofurantoin resistance. Nitrofurantoin-susceptibility testing showed that the algorithm correctly predicted susceptibility levels for double mutants with known resistance-associated genetic alterations and for single mutants with novel genetic alterations (Table 4). The discrepancy between predicted susceptibility and MICs of isolates carrying single resistance-associated alterations (which include confirmed loss-of-function alterations, such as nonsense and frameshift mutations, gene interruptions and truncations) in *nfsA* or *nfsB* suggests that impaired function or production of both oxygen-insensitive nitroreductases NfsA and NfsB are required to render *E. coli* resistant to nitrofurantoin. This discrepancy also implies that we may have overestimated the prevalence of nitrofurantoin resistance in the sequenced collection of UK *E. coli* isolates, although the estimate of 6.3% is close to the observed prevalence of 5% in England by 2019 (10, 16, 17).

To provide reference information for future research, we have developed database NITREc (github.com/wanyuac/NITREc), which will facilitate searching for the most closely related reference sequence of a query allele of *nfsA*, *nfsB*, or *ribE*, minimising the number of reported variants and reducing noise in the results. We anticipate improvement in accuracy of both variant identification and susceptibility prediction when new references are incorporated into the database in the future.

This study has several limitations. First, the impact of novel missense mutations in *nfsA* and *nfsB* on nitroreductase function and bacterial fitness were not experimentally confirmed, for example, through mutagenesis experiments. Such confirmation is desirable as the observed variation in MICs of single mutants carrying missense mutations indicates that PROVEAN-predicted resistance mutations may not cause reduced nitrofurantoin susceptibility and their effects may be offset by other susceptibility mechanisms. Second, epidemiological analysis of nitrofurantoin-resistant *E. coli* is limited owing to a lack of available information about nitrofurantoin susceptibility of *E. coli* isolates in the UK or abroad. As such, several key clinical questions have not been addressed in this study, including the association of STs or infection sites (e.g., blood, urinary tract, etc.) with nitrofurantoin resistance, co- or cross-resistance between nitrofurantoin and other antimicrobials, and temporal trend of nitrofurantoin resistance in the UK. Third, regulatory elements of the nitroreductase system or DNA-repair mechanisms were not investigated in this study. Furthermore, untargeted analysis was not carried out for identifying novel genetic mechanisms of nitrofurantoin resistance due to the small sample size of nitrofurantoin-resistant *E. coli* isolates that were available in this study.

## Conclusions

We predict the major cause of nitrofurantoin resistance in UK *E. coli* to comprise sporadic *de novo* mutations in chromosomal genes *nfsA*, *nfsB*, and *ribE*; and interruptions of *nfsA* and *nfsB* by insertion sequences. Accordingly, clonal expansion of resistant mutants, mobilisation of insertion sequences, and selection of resistant clones in the presence of nitrofurantoin are believed to be three driving forces in the evolution of nitrofurantoin-resistant *E. coli* in the UK. Previous reports have suggested that *nfsA* and *nfsB* mutations are associated with significant fitness cost that may obstruct propagation of resistant *E. coli* (35, 48). However, it is notable that isolates investigated in these reports and the current research were all identified from clinically significant infections, implying a fitness to survive in enteric microbiota and to survive host immune response in the lower urinary tract. This fitness was also supported by the possible community transmission identified with clonal isolates IN01 and IN02. The same nitrofurantoin-resistance determinants of IN01–09 were also identified in bloodstream isolates, underlining the ability of these *E. coli* to adapt to circumstance.

It is unclear whether nitrofurantoin resistance will become more prevalent as a result of increased community nitrofurantoin exposure following the change in the national guideline. As such, routine nitrofurantoin susceptibility testing and WGS of *E. coli* isolates are needed to monitor this trend. The tools provided in our study will facilitate future WGS-based surveillance. Further work is required to assess the fitness cost of resistance-associated genetic alterations and to identify novel mechanisms of nitrofurantoin resistance.

## Materials and Methods

### *E. coli* isolates and antimicrobial susceptibility testing

Automated EUCAST disc diffusion tests for nitrofurantoin susceptibility (Oxoid™ CT0034B 100-μg nitrofurantoin discs, ThermoFisher Scientific™, USA) in clinical urinary *E. coli* isolates were routinely performed by a microbiology laboratory of the Imperial College Healthcare NHS Trust in north west London. The laboratory served a population of 2.5 million. The research use of anonymised, residual samples obtained for routine diagnostic purposes was approved by a research ethics committee (IRAS Project ID 162013; REC reference 06/Q0406/20). In the period 2018–2019, 18 nitrofurantoin-resistant *E. coli* isolates were collected. Nonetheless, the susceptibility result could be reproduced for only nine of these isolates (IN01–09) when their frozen stock was retrieved and then retested using the same disc diffusion method. Aerobic MIC of nitrofurantoin was measured for each of the nine isolates using MIC Test Strips (Liofilchem^®^, Italy) with nitrofurantoin concentrations ranging between 0.032–512 mg/L. Nitrofurantoin-susceptible *E. coli* isolate EC0098B (nitrofurantoin MIC: 32 mg/L) was chosen as a negative control. MICs were reported as per conventional two-fold series of concentration increments (2, 4, 8, … mg/L) and interpreted in accordance with EUCAST breakpoints v10.0 (49).

In order to identify genetic variants related to nitrofurantoin resistance using a comparative approach, the NCBI Pathogen Detection portal (www.ncbi.nlm.nih.gov/pathogens/), NCTC 3000 Project, EnteroBase, NIHR HPRU isolate collection (BioProject accession: PRJEB20357; manuscript in preparation), and literature were searched for *E. coli* genomes from isolates with known nitrofurantoin susceptibility profiles (7, 33, 34, 50, 51). Complete and draft genome assemblies were downloaded from the NCBI nucleotide database and sequence reads of unassembled genomes were downloaded from NCBI Sequence Read Archive (SRA). In total, 208 *E. coli* genomes were obtained from the search, consisting of 196 genomes downloaded from the GenBank database (www.ncbi.nlm.nih.gov/genbank/) and 12 genomes from clinical isolates collected from north west London by the HPRU in 2015–2016.

### Whole-genome DNA extraction and sequencing

Genomic DNA of isolates IN01–09 was extracted using a phenol/chloroform method (52), checked for integrity using gel electrophoresis, and stored at −20°C. DNA libraries were prepared in a paired-end layout with NEBNext^®^ Ultra™ II FS DNA Library Prep Kit (New England BioLabs, UK) and sequenced under a 150-bp read length using Illumina HiSeq 4000 systems and HiSeq SBS Kit v4 reagents (Illumina, USA). Read demultiplexing and adapter trimming were performed by the Imperial BRC Genomics Facility using their production pipeline.

### Quality control of sequence reads

Quality summaries of all read files were generated using FastQC v0.11.9 and compiled using MultiQC v1.8 (53, 54). With Trimmomatic v0.39, the reads of IN01–09 were trimmed for an average base quality of Phred Q20 in a 5-bp sliding window and were then filtered for a minimum length of 50 bp (55). Seqkit v0.12.0 was used for randomly sampling the filtered reads of each genome to an 80-fold coverage, assuming an average genome size of 5 Mbp (56, 57). For reads downloaded from the SRA, case-by-case read trimming and filtering was conducted using Trimmomatic in order to deal with large variation in read quality and to obtain a coverage of less than 100 folds. Genomes having less than 40-fold coverages were excluded from further analysis. DNA contamination in reads was evaluated for each genome using Kraken v2.0.8-beta and its full bacterial database (accessed in February 2020) (58).

### *De novo* genome assembly and annotation

Reads of each genome were assembled using Unicycler v0.4.9b (59). Quality of assemblies were evaluated based on summary statistics calculated by QUAST v5.0.2 (60). In summary, 110–236 contigs (median: 137) and an assembly length of 4.6–5.3 Mbp (median: 4.9 Mbp) were obtained per genome (Table S2). Complete plasmid sequences were identified in Bandage visualisation of assembly graphs (61). Assemblies were annotated through Prokka v1.13 with a reference protein database representing proteins extracted from all *E. coli* genomes that were publicly available in the NCBI RefSeq database by March 2020 (62). Redundant protein sequences were removed from this database using CD-HIT v4.8.1 (63) under a minimum amino acid identity of 70%.

### Estimating population structure

Multi-locus sequence typing (MLST) of genomes IN01–09 was conducted using ARIBA v2.14.4 and the MLST of other collected genomes were conducted using mlst (github.com/tseemann/mlst) (64). The Achtman scheme for *E. coli*, which was downloaded from PubMLST database (pubmlst.org) with ARIBA in March 2020, was used for the MLST analysis. Core-genome population structure was estimated through clustering all genome assemblies with PopPUNK v2.0.2 given k-mer lengths increased from 15 bp to 23 bp by a 2-bp step size (23).

### Identifying genetic alterations of *nfsA*, *nfsB*, and *ribE*

For each of sample genomes IN01–09, the most closely related complete genome of a nitrofurantoin-susceptible isolate was chosen from our collection as a reference according to core-genome distances calculated by PopPUNK. Reads of the sample genome were then mapped to the chromosome sequence of this reference genome using Bowtie v2.4.1 under the mode sensitive-local (65). Single-nucleotide variants (SNVs) and indels were identified from mapped reads using BCFtools v1.9 when filtering out low-confidence variant calls (QUAL ≤20, DP ≤10, or MQ ≤20) as well as low-quality base calls (Phred Q <20). Moreover, SNVs and indels within repetitive regions identified in the reference sequence with nucmer v3.1 were excluded for accuracy (66). All remaining variants in the sample genome were annotated using SnpEff v4.3t and a database built from the reference sequence (67).

Bandage v0.8.1 (61) was used for searching for loci of *nfsA*, *nfsB*, and *ribE* in genome assemblies of IN01–09 and extracting allele sequences. SNVs and indels were identified in these alleles using web-based megaBLAST (blast.ncbi.nlm.nih.gov) and compared to those identified through read mapping. Interruptive insertion sequences were inferred from assembly graphs using Bandage and were searched against the ISFinder database for reference sequences and classification (68). Insertion sites of these putative insertion sequences were determined with ISMapper v2.0.1 (69). Copy numbers of *nfsA*, *nfsB*, and *ribE* in each sample genome were predicted from mapped reads using CNOGpro v1.1 (100-bp sliding windows and 1,000 bootstrap samples per gene) (70, 71). Translated sequences of these three genes were extracted from Prokka annotations and were aligned using the ClustalW algorithm implemented in MEGA X (72, 73). Functional effect of each amino acid substitution was predicted using PROVEAN Protein (accessed in June 2020).

### Validating interrupted *nfsA* and *nfsB*

PCR experiments were conducted to confirm insertion sequences that might have interrupted *nfsA* and *nfsB* in genomes IN01–03. Plausible sequence of each interrupted region was reconstructed from the assembly graph using Bandage and thereby 22-bp PCR primers (Table 5) were designed using Primer3web v4.1.0 (primer3.ut.ee) (74). Primer positions in template genomic regions and expected PCR products are illustrated in Fig. 2. PCR was carried out using a T100™ Thermal Cycler (Bio-Rad, USA) and GoTaq^®^ DNA Polymerase (Promega, UK). PCR primers and products were sent to Genewiz (UK) for Sanger sequencing. Reads were trimmed for removing ambiguous bases before alignment.

**TABLE 5.**
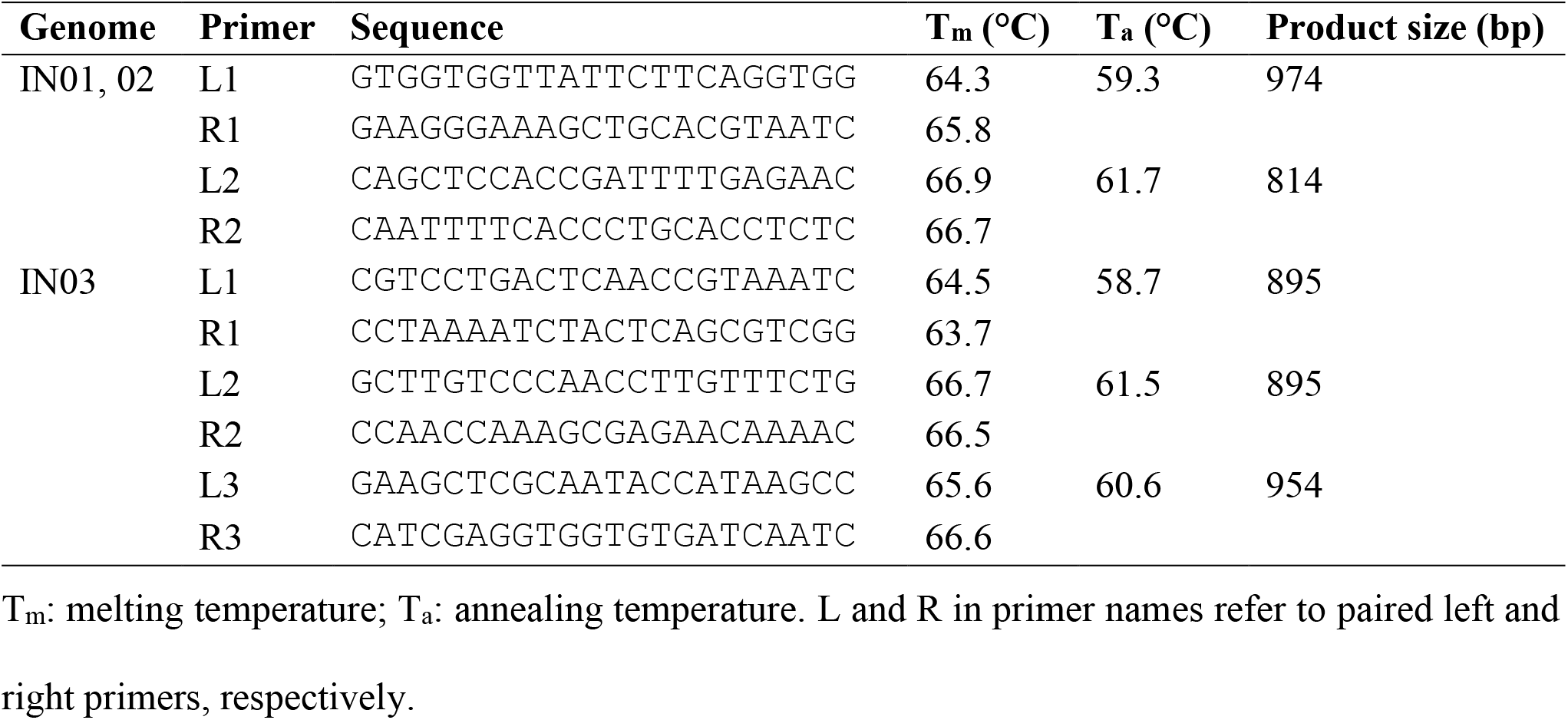
PCR primers for validating interrupted genetic regions in genome IN01–03.

### Detecting acquired antimicrobial resistance genes

An ARIBA-compatible reference database of acquired AMR genes was created from the ResFinder database (commit hash: 7e1135b) through quality filtering and sequence clustering (nucleotide identity ≥80%) using ARIBA and CD-HIT-EST v4.8.1 (29, 63). Acquired AMR genes in genomes IN01–09 were detected from sequence reads using ARIBA and this reference database, whereas these genes in other *E. coli* genomes were identified from genome assemblies using ABRicate (github.com/tseemann/abricate) (75) and the ResFinder database without clustering.

### Screening for nitrofurantoin-resistance determinants in the UK *E. coli* population

Genome assemblies of UK *E. coli* isolates were downloaded from NCBI nucleotide databases (as of August 2020; Table S3) or retrieved from our ongoing NIHR HPRU study of *E. coli* bacteraemia and intestinal colonisation. Collection dates and locations of isolates were retrieved from the NCBI BioSample database, NCTC Bacteria and Mycoplasmas Browse (www.phe-culturecollections.org.uk), EnteroBase, and related literature. This information was manually inspected for accuracy and non-redundancy. A non-redundant reference database NITREc was created from allelic and translated sequences of *nfsA*, *nfsB*, and *ribE* in all the 217 *E. coli* isolates of known nitrofurantoin susceptibility (Fig. 1). Specifically, CD-HIT-EST and CD-HIT were used for sequence deduplication. Computer scripts developed for gene detection and mutation identification are available in the NITREc code repository (github.com/wanyuac/NITREc/tree/master/Script) (including those mentioned below).

Nucleotide BLAST (megaBLAST) v2.9.0 (76) was used for identifying *nfsA*, *nfsB*, *ribE*, *oqxA*, and *oqxB* and for extracting their allele sequences (script screenGenes.pbs), which were then translated into protein sequences using script translateDNA.py. Particularly, alleles of *nfsA*, *nfsB*, and *ribE* from nitrofurantoin susceptible *E. coli* strain ATCC25922 and alleles of *oqxA* and *oqxB* from the ResFinder database were used as queries for gene screen. Since missense mutations in the start or stop codon may reduce the length of a BLAST alignment, hits showing partial or complete truncation of either codon were manually verified. CD-HIT-EST and CD-HIT were used for identifying identical nucleotide and protein sequences, respectively. Sequence alignments were generated using Clustal Omega (www.ebi.ac.uk/Tools/msa/clustalo). Nonsense mutations were determined through comparing lengths of predicted protein sequences to those of their reference sequences. Missense mutations were identified in the alignments of protein sequences using script missenseFinder.py, which identifies amino acid substitutions of each query protein sequence by comparing it to its most closely related reference protein sequence in the NITREc database. Finally, using script findKnownMutations.py, missense mutations were searched for those known to be associated with nitrofurantoin resistance (Tables A1–3, and Table 1).

### Evolutionary analysis of missense mutations in *nfsA*, *nfsB*, and *ribE*

An ML estimate of the ratio *ω* of each gene was estimated from the codon alignment using a reference-free approach implemented in GenomegaMap v1.0.1 (77), assuming a constant *ω* per gene. Specifically, for each gene, full-length, indel-free alleles were translated into protein sequences using NITREc script translateDNA.py (codon table: 11). Premature proteins (due to nonsense mutations) and their corresponding alleles were identified and then excluded using NITREc script rmProteinsByLength.py. A multi-sequence alignment of remaining alleles was generated using program MUSCLE in software package MEGA X (78) and was then converted into a codon alignment using script pal2nal.pl (79).

For each of genes *nfsA*, *nfsB*, and *ribE*, SNP sites were identified in a deduplicated set of allele sequences that were used for estimating the ω ratios — namely, untruncated alleles that only carried missense mutations. A maximum-likelihood (ML) phylogenetic tree was reconstructed for each gene from the alignment of deduplicated alleles. Specifically, nucleotide sequences were deduplicated using CD-HIT-EST before tree reconstruction. An extended selection of substitution models by IQ-Tree v1.6.12 (parameters: −t BIONJ −m MF −mtree) was performed for each gene (80). Since the best-fit model differed between genes, candidate models of each gene were sorted in an ascending order of models’ Bayesian information criterion (BIC) scores, and the model consistently ranked < 10 across all three genes was chosen for the tree reconstruction. Then an ML tree was reconstructed for each gene by IQ-Tree with the chosen model (TIM3e + R2), a BIONJ starting tree, 10 independent runs (for selecting the tree of the greatest maximum likelihood), and 500 bootstrap replicates (for supporting branches in the selected tree).

Identification of variant sites in the sequence alignment used for reconstructing each gene tree was performed with snp-sites v2.5.1 (81). Homoplasy sites in the alignment were identified and plotted using R package homoplasyFinder (82), which took as input the gene tree in addition to the allele alignment. A PHI was calculated from the allele alignment of each gene and the significance of PHI was tested for using PHIPack (83) with a 80-bp sliding window, given the null hypothesis of no recombination. Recombination within a gene was considered significant if the p-value of a PHI did not exceed 0.05.

### Predicting nitrofurantoin susceptibility from detected genetic diversity

The prediction of nitrofurantoin susceptibility was conducted independently for each HPRU isolate at sequence, genetic, and genomic levels. It also considered whether the gene of interest is intrinsic or acquired. The prediction method can be concisely described using the following scoring algorithm, which assumes that loss of intrinsic gene function and gain of both functional *oqxA* and *oqxB* alleles are associated with reduced nitrofurantoin susceptibility. Calculation of gene-level and genome-level scores have been implemented in NITREc helper script scoreHitsNITR.R, whereas manual inspections are required to determine sequence-level scores.

Firstly, at the sequence level, supposing *n_i_* BLAST hits of gene *i* (*i = nfsA, nfsB, ribE, oqxA, or oqxB*) are identified in a genome, a score *s_ij_* (*j* = *1, …, n_i_*) is assigned to each hit to represent the probability of this sequence to confer reduced nitrofurantoin susceptibility. Then an arbitrary value is assigned to the score for each hit of intrinsic genes: *s_ij_* = *1*, if the sequence is truncated or interrupted (which cannot be distinguished from each other by BLAST without assembly graphs) or has a start-codon loss, a frameshift or nonsense mutation, or any missense mutation known to be associated with nitrofurantoin resistance; *s_ij_* = *0.1*, if the sequence only has missense mutations whose impact on nitrofurantoin susceptibility remains unknown; and *s_ij_* = 0, if the sequence is identical to any known “wildtype” allele present in a nitrofurantoin-susceptible isolate. On the other hand, for each hit of acquired genes, *s_ij_* = *1*, if the sequence is identical to a reference allele in the ResFinder database; *s_ij_* = *0.1*, if the sequence only carries missense mutations of unknown impacts on the protein function; and *s_ij_* = *0*, if the sequence is truncated or interrupted (No other kind of *oqxA* or *oqxB* variants were found in this study).

Secondly, a gene score *g_i_* (*i* = *nfsA, nfsB, ribE, oqxA, oqxB*) is determined from sequence scores. Specifically, for each intrinsic gene, *g_i_* = *min{s_ij_}*, where *j* = *1, …, n_i_*. Namely, gene *i* is believed to not confer reduced nitrofurantoin susceptibility when there is at least one wildtype allele. For acquired genes *oqxA* and *oqxB*, however, *g_i_* = *max{s_ij_}* (*j* = *1, …, n_i_*) as wildtype alleles are believed to confer reduced nitrofurantoin susceptibility. Moreover, *g_i_* = *1* and *g_i_*’ = *0* when intrinsic gene *i* and acquired gene *i*’ are not detected, respectively.

Finally, a risk score *r* is calculated from gene scores for each genome. Let *g_1_*, …, *g_5_* represent gene scores of *nfsA*, *nfsB*, *ribE*, *oqxA*, and *oqxB*, respectively, then each gene score takes a value of 0, 0.1, or 1. Since the multidrug effulx pump OqxAB requires both functional components OqxA and OqxB to reduce the intracellular concentration of nitrofurantoin, the risk score is calculated by formula:

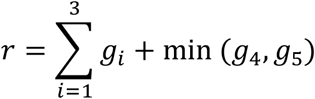

Hence *r* takes values between zero (nitrofurantoin susceptible) and four (nitrofurantoin resistant). In our study, genomes were sorted by risk scores to identify plausible resistant and susceptible isolates. Predicted nitrofurantoin susceptibility was labelled and interpreted in accordance with the EUCAST terminology: S (susceptible, standard dosing regimen), if *r* = *0*; I (susceptible, increased exposure), if *r* = *1*; R (resistant), if *r* ≥ *2*; S/I, if *r* = *0.1*; and S/I/R, if *0.1* < *r* < *1* or *1* < *r* < *2*.

### Validating predictions of nitrofurantoin resistance

Since HPRU isolates of known nitrofurantoin susceptibility (based on clinical records) had been used by the prediction algorithm as references, we retrieved HPRU isolates of unknown nitrofurantoin susceptibility for validating the predictions. Nitrofurantoin MICs were determined using Etests (Liofilchem^®^ MIC Test Strips). First, 20 isolates that were anticipated to show reduced nitrofurantoin susceptibility were selected for Etests (Tables 3 and 4). Second, additional seven HPRU isolates were included for Etests because these isolates had mutations detected in *nfsA* or *nfsB* of IN08, or carried an alternative stop codon or novel mutation in *ribE*, although the predicted susceptibility was uncertain due to a lack of information from literature (Table 4). Furthermore, disc diffusion tests (as per EUCAST manual v8.0) (84) and MALDI-TOF mass spectrometry were conducted for confirming the heterogeneous responses to nitrofurantoin. A positive control (IN09) and a negative control (ATCC25922) were included for experiments. (Doc. S1)

## Data availability

Sequence reads and genome assemblies of isolates IN01–09 have been deposited in the European Nucleotide Archive (ENA) under BioProject accession number PRJEB38850. Nucleotide and protein sequences of *nfsA, nfsB*, and *ribE* alleles in genomes of these nine isolates can be accessed in our database NITREc. The database also offers nucleotide sequences of genetic elements related to the interrupted *nfsA/nfsB* regions in IN01–03 (github.com/wanyuac/NITREc/tree/master/Seq/Nucl/IS). All supplementary files are available on figshare (doi.org/10.6084/m9.figshare.14632653.v1).

## Supplementary materials

TABLE S1, Excel file. Sample information and ENA accessions of *E. coli* isolates IN01–09. Bacterial susceptibility to trimethoprim, ciprofloxacin, gentamicin, co-amoxiclav, cephalexin, amoxicillin, amikacin, meropenem, cefoxitin, ceftazidime, piperacillin-tazobactam, cefotaxime, mecillinam, ceftazidime-avibactam, ertapenem, temocillin, and fosfomycin was automatically tested for by the same hospital microbiology laboratory using the EUCAST disc diffusion method.

TABLE S2, Excel file. Supplementary results from analysing genomes of isolates IN01–09.

TABLE S3, Excel file. Accessions and sample information of public genome assemblies used for screening nitrofurantoin-resistance determinants in the UK *E. coli* population.

TABLE S4, Excel file. Amino acid substitutions identified in NfsA, NfsB, and RibE of 12,412 UK *E. coli* genomes.

TABLE S5, Excel file. Genotypes and predicted nitrofurantoin susceptibility of 69 HPRU *E. coli* isolates that were used for the validation/exploratory experiment.

FIG S1, PDF image. Homoplasy detected in *nfsA* alleles by homoplasyFinder. Left: an ML, midpoint-rooted, and ladderised phylogenetic tree of 328 *nfsA* alleles (723 bp, no nonsense mutation) identified in 12,100 BLAST hits of *nfsA* (Fig. 3) in our collection of UK *E. coli* genomes. Tips of the tree are labelled by names of representative genomes where the alleles were present. Internal nodes are labelled by homoplasy-SNP positions deduced through ancestral reconstruction. Right: Genotype matrix of 82 homoplasy SNP sites in these *nfsA* alleles. Positions of these sites are labelled at the top of the matrix.

FIG S2, PDF image. Homoplasy detected in representative *nfsB* alleles by homoplasyFinder. Left: an ML, midpoint-rooted, and ladderised phylogenetic tree of 231 *nfsB* alleles (654 bp, no nonsense mutation) identified in 12,319 BLAST hits of *nfsB* (Fig. 3) in our collection of UK *E. coli* genomes. Tips of the tree are labelled by names of representative genomes where the alleles were present. Internal nodes are labelled by homoplasy-SNP positions deduced through ancestral reconstruction. Right: Genotype matrix of 50 homoplasy SNP sites in these *nfsB* alleles. Positions of these sites are labelled at the top of the matrix.

FIG S3, PDF image. Homoplasy detected in representative *ribE* alleles by homoplasyFinder. Left: an ML, midpoint-rooted, and ladderised phylogenetic tree of 88 *ribE* alleles (471 bp, no nonsense mutation) identified in 12,407 BLAST hits of *ribE* (Fig. 3) in our collection of UK *E. coli* genomes. Tips of the tree are labelled by names of representative genomes where the alleles were present. Internal nodes are labelled by homoplasy-SNP positions deduced through ancestral reconstruction. Right: Genotype matrix of 12 homoplasy SNP sites in these *ribE* alleles. Positions of these sites are labelled at the top of the matrix.

DATASET S1, 7-Zip compressed directory consisting of assembly graphs of genomes IN01–09 and the ML phylogenetic trees of *nfsA*, *nfsB*, and *ribE*.

DOC S1, PDF document. Methods for the experimental validation of predicted nitrofurantoin susceptibility and the identification of nitrofurantoin-resistant subpopulations of heterogeneous *E. coli* isolates.

## Acknowledgments

The authors acknowledge the support of the National Institute for Health Research (NIHR) Imperial College Biomedical Research Centre (BRC), which funds the BRC Infection Bioresource, the Imperial BRC Genomics Facility, and the Colebrook Laboratory. The Imperial BRC Genomics Facility has provided resources and support that have contributed to the research results reported within this paper. We are also grateful to the staff in the Imperial College NHS Healthcare Trust Diagnostic Laboratory for their generous support and expertise.

We thank Ho Kwong Li, Kristin Krohn Huse, Max Pearson, Matthew K. Siggins, Hanqi Li (Imperial College London), and Rūta Prakapaitė (the University of Basel) for sharing experimental expertise. We express our gratitude to John A. Lees (Imperial College London) for advice on using software PopPUNK.

EJ is a Rosetrees/Stoneygrate 2017 Imperial College Research Fellow, funded by Rosetrees Trust and the Stoneygate Trust (Reference number: M683). NJC was supported by a Sir Henry Dale Fellowship, jointly funded by the Wellcome Trust and Royal Society (104169/Z/14/A), and a Medical Research Council (MRC) grant (MR/T016434/1). YW was partly supported by the NIHR Senior Investigator Award of Professor Alison Holmes (Imperial College London) and by the Imperial College Biomedical Research Centre.

## Authors’ contributions

Experimental design, primary genomic analysis, prediction algorithm: YW; Laboratory work: EM, RCYL, XZ, YW; Comparative genomics, evolutionary analyses, and interpretation: AV, EJ, NJC, and YW; Project conception: SS, MJE, and NW; Project supervision: SS and MJE; First draft of the manuscript: YW; Manuscript revisions: SS, EJ, AV, NJC, MJE, NW, and RCYL; Final manuscript editing and approval: all authors.

## Appendices

In this section, we list known missense mutations in NfsA, NfsB, and RibE that were exclusively identified in *E. coli* isolates of reduced nitrofurantoin susceptibility. Particularly, we only consider experimentally confirmed (such as using stepwise mutations) causative mutations and those having a deleterious impact on protein structure or translation by prediction. Absence of these mutations in known nitrofuranoin-susceptible *E. coli* was confirmed through aligning wildtype protein sequences from the NITREc database with MEGA X.

**TABLE A1.** NfsA missense mutations associated with reduced nitrofurantoin susceptibility in *E. coli*.

| Pos. | Ref. | Alt. | Ref. accession | Alt. accession | Confidence | Citation |
| --- | --- | --- | --- | --- | --- | --- |
| 1 | M | I | NG | NA | Possible reduced translation | (35) |
| 11 | H | Y | WP_000189141.1<sup>+</sup> | QEL51189.1 | PROVEAN prediction | (36) |
| 11 | H | Y | AAC73938.1 | WP_119187568.1<sup>§</sup> | PROVEAN prediction | (85) |
| 11 | H | L | AAC73938.1<sup>*</sup> | NA | Experimentally confirmed | (6) |
| 15 | R | C | NG | NA | Experimentally confirmed | (35, 41) |
| 33 | S | R | AAC73938.1 | WP_094323839.1<sup>§</sup> | PROVEAN prediction | (35, 85) |
| 33 | S | N | WP_000189165.1<sup>+</sup> | QEL51188.1 | PROVEAN prediction | (36) |
| 36 | A | V | NG | NA | PROVEAN prediction | (41) |
| 38 | S | Y | WP_000189167.1<sup>+</sup> | QEL51184.1 | PROVEAN prediction | (36) |
| 67 | Q | L | AAC73938.1 | NA | PROVEAN prediction | (85) |
| 80 | C | R | AAC73938.1 | NA | PROVEAN prediction | (85) |
| 89 | I | N | AAC73938.1<sup>*</sup> | NA | Experimentally confirmed | (6) |
| 126 | G | R | AAC73938.1 | WP_023149811.1<sup>\$</sup> | PROVEAN prediction | (85) |
| 131 | G | D | NG | NA | Previously reported | (35) |
| 133 | R | S | AAC73938.1<sup>#</sup> | WP_096948202.1 | In active-site pocket | (20) |
| 154 | G | E | AAC73938.1 | WP_000189151.1<sup>\$</sup> | PROVEAN prediction | (85) |
| 158 | G | C | NG | NA | PROVEAN prediction | (41) |
| 203 | R | L/A | NG | NA | Experimentally confirmed | (35, 37) |
| 203 | R | C | AAC73938.1<sup>#</sup> | WP_072644388.1 | Experimentally confirmed | (20, 35, 37, 85) |
| **212** | **W** | **R** | WP_000189134.1<sup>+</sup> | QEL51185.1 | PROVEAN prediction | (36) |

**TABLE A2.** NfsB missense mutations associated with reduced nitrofurantoin susceptibility in *E. coli*.

| Pos. | Ref. | Alt. | Ref. accession | Alt. accession | Confidence | Citation |
| --- | --- | --- | --- | --- | --- | --- |
| 1 | M | I | NG | NA | Possible reduced translation | (35) |
| 33 | L | P | AAC73679.1<sup>*</sup> | NA | Experimentally confirmed | (6) |
| 34 | L | R | AAC73679.1<sup>*</sup> | WP_096943843.1 | Experimentally confirmed | (6) |
| 37 | S | R | AAC73679.1<sup>*</sup> | NA | Experimentally confirmed | (6) |
| 40 | S | C | AAC73679.1<sup>*</sup> | NA | Experimentally confirmed | (6) |
| 44 | Q | L | NG | NA | Experimentally confirmed | (35) |
| 44 | Q | P | AAC73679.1<sup>*</sup> | NA | Experimentally confirmed | (6) |
| 44 | Q | H | AAC73679.1 | EFN8391251.1<sup>\$</sup> | PROVEAN prediction | (85) |
| 45 | P | S | WP_000351450.1<sup>+</sup> | QEL51196.1 | PROVEAN prediction | (36) |
| 45 | P | Q | AAC73679.1* | NA | Experimentally confirmed | (6) |
| 71 | N | K | AAC73679.1* | NA | Experimentally confirmed | (6) |
| 84 | F | S | WP_000351450.1<sup>+</sup> | QEL51192.1 | PROVEAN prediction | (36) |
| 84 | F | S | AAC73679.1 | NA | PROVEAN prediction | (85) |
| 93 | V | G | NG | NA | Experimentally confirmed | (35) |
| 105 | D | Y | AAC73679.1* | NA | Experimentally confirmed | (6) |
| 107 | R | H | NG | NA | Experimentally confirmed | (35) |
| 107 | R | C | AAC73679.1 | WP_096996177.1<sup>§</sup> | PROVEAN prediction | (85) |
| 138 | W | C | NG | NA | Experimentally confirmed | (35) |
| 148 | G | S | AAC73679.1* | NA | Experimentally confirmed | (6) |
| 165 | E | A | NG | NA | Experimentally confirmed | (35) |
| 166 | G | D | NG | NA | Experimentally confirmed | (35) |
| **192** | **G** | **D/A** | NG | NA | Experimentally confirmed | (35) |
| 192 | G | S | AAC73679.1 | NA | PROVEAN prediction | (85) |
| 207 | R | H | AAC73679.1* | NA | Experimentally confirmed | (6, 85) |

**TABLE A3.**
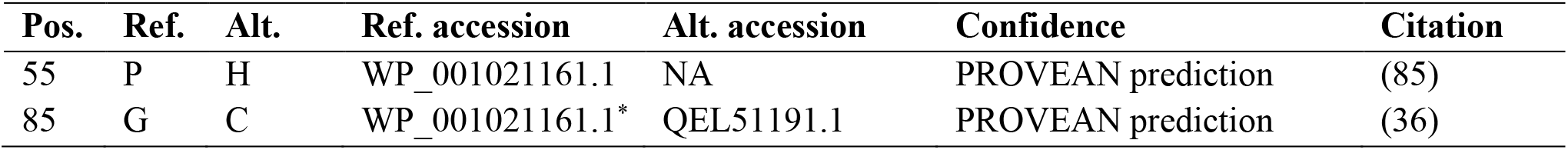
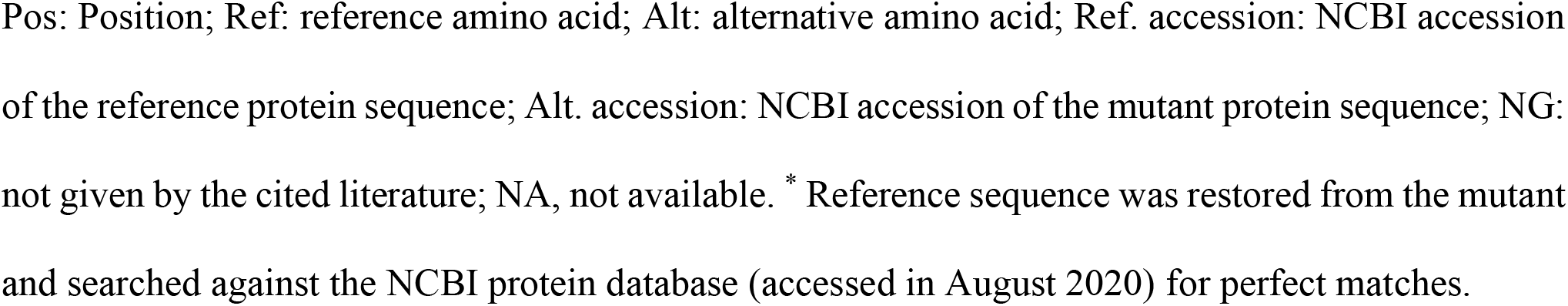
RibE missense mutations associated with reduced nitrofurantoin susceptibility in *E. coli*.

| Pos. | Ref. | Alt. | Ref. accession | Alt. accession | Confidence | Citation |
| --- | --- | --- | --- | --- | --- | --- |
| 55 | P | H | WP_001021161.1 | NA | PROVEAN prediction | (85) |
| 85 | G | C | WP_001021161.1* | QEL51191.1 | PROVEAN prediction | (36) |

Pos: Position; Ref: reference amino acid; Alt: alternative amino acid; Ref. accession: NCBI accession of the reference protein sequence; Alt. accession: NCBI accession of the mutant protein sequence; NG: not given by the cited literature; NA, not available. \* Reference sequence was restored from the mutant and searched against the NCBI protein database (accessed in August 2020) for perfect matches.

